# Pharmacologic activation of the mitochondrial phospho*enol*pyruvate cycle enhances islet function in vivo

**DOI:** 10.1101/2020.02.13.947630

**Authors:** Abudukadier Abulizi, Romana Stark, Rebecca L. Cardone, Sophie L. Lewandowski, Xiaojian Zhao, Tiago C. Alves, Craig Thomas, Charles Kung, Bei Wang, Stephan Siebel, Zane B. Andrews, Matthew J. Merrins, Richard G. Kibbey

## Abstract

The mitochondrial GTP (mtGTP)-dependent phospho*enol*pyruvate (PEP) cycle is an anaplerotic-cataplerotic mitochondrial shuttle utilizing mitochondrial PEPCK (PCK2) and pyruvate kinase (PK). PEP cycling stimulates insulin secretion via OxPhos-independent lowering of ADP by PK. We assess *in vivo* whether islet PCK2 is necessary for glucose sensing and if speeding the PEP cycle via pharmacological PK activators amplifies insulin secretion. *Pck2*^*-/-*^ mice had severely impaired insulin secretion during islet perifusion, oral glucose tolerance tests and hyperglycemic clamps. Acute and chronic pharmacologic PK activator therapy improved islet insulin secretion from normal, high-fat diet (HFD) fed, or Zucker diabetic fatty (ZDF) rats, and glucolipotoxic or diabetic humans. A similar improvement in insulin secretion was observed in regular chow and HFD rats *in vivo*. Insulin secretion and cytosolic Ca^2+^ during PK activation were dependent on PCK2. These data provide a preclinical rationale for strategies, such as PK activation, that target the PEP cycle to improve glucose homeostasis.

**Highlights:**

- Loss of mitochondrial phospho*enol*pyruvate (PEP) impairs insulin release *in vivo*.
- Pyruvate kinase (PK) activators stimulate beta-cells in preclinical diabetes models.
- PEP cycling *in vivo* depends on PK and mitochondrial PEPCK (PCK2) for insulin release.
- Acute and 3-week oral PK activator amplifies insulin release during hyperglycemia.

**eTOC Blurb:** Abudukadier et al. show that small molecule pyruvate kinase activation *in vivo* and *in vitro* increases insulin secretion in rodent and human models of diabetes. The phospho*enol*pyruvate (PEP) cycling mechanism and its amplification are dependent on mitochondrial PEPCK (PCK2).

## Introduction

A foregone conclusion from the consensus model of beta-cell glucose sensing is that glucokinase (GK)-regulated OxPhos lowers ADP to control plasma membrane depolarization associated with insulin secretion. Pharmacological targeting of either glucose entry at the beginning (GK activators) or depolarization at the end (sulphonylureas) of this canonical pathway stimulate secretion, but have had incrementally diminished long term therapeutic success (De Ceuninck et al., 2013; Efanova et al., 1998; Erion et al., 2014; Iwakura et al., 2000). Successful targeting of the intermediate steps in the canonical pathway by other approaches, including enhanced OxPhos, have also been of limited success with no agents currently approved for use in humans. The OxPhos-independent mitochondrial GTP (mtGTP)-dependent phospho*enol*pyruvate (PEP) cycle is a metabolic signaling pathway important for nutrient-stimulated insulin secretion that provides an opportunity as a therapeutic target (**Figure 1A**). This pathway begins with the entry of glucose-derived pyruvate into the TCA cycle (anaplerosis) via the mitochondrial enzyme pyruvate carboxylase (PC) to generate oxaloacetic acid (OAA). OAA exits from the TCA cycle (cataplerosis) as PEP following the action of mtGTP-dependent PCK2 (Alves et al., 2015; Jesinkey et al., 2019; Kibbey et al., 2007; Stark et al., 2009). The cycle is completed when the PEP is then enzymatically hydrolyzed back to pyruvate by pyruvate kinase (PK) in order to lower cytosolic ADP. PK-mediated ADP lowering first slows down OxPhos and then closes K_ATP_ channels to depolarize the plasma membrane (Lewandowski et al., 2019). There are also, cell-permeable, small-molecule allosteric PK activators that lower the K_m_ for PEP, increase the frequency of metabolic and electrical oscillations and stimulate insulin release. As a potential therapeutic strategy *not predicted by the consensus model*, we question whether accelerating the PEP cycle with intravenous or oral delivery of PK activators 1) works *in vivo*, 2) is dependent on PCK2, and 3) can enhance insulin secretion in preclinical rodent models and diabetic human islets.

**Figure 1.**
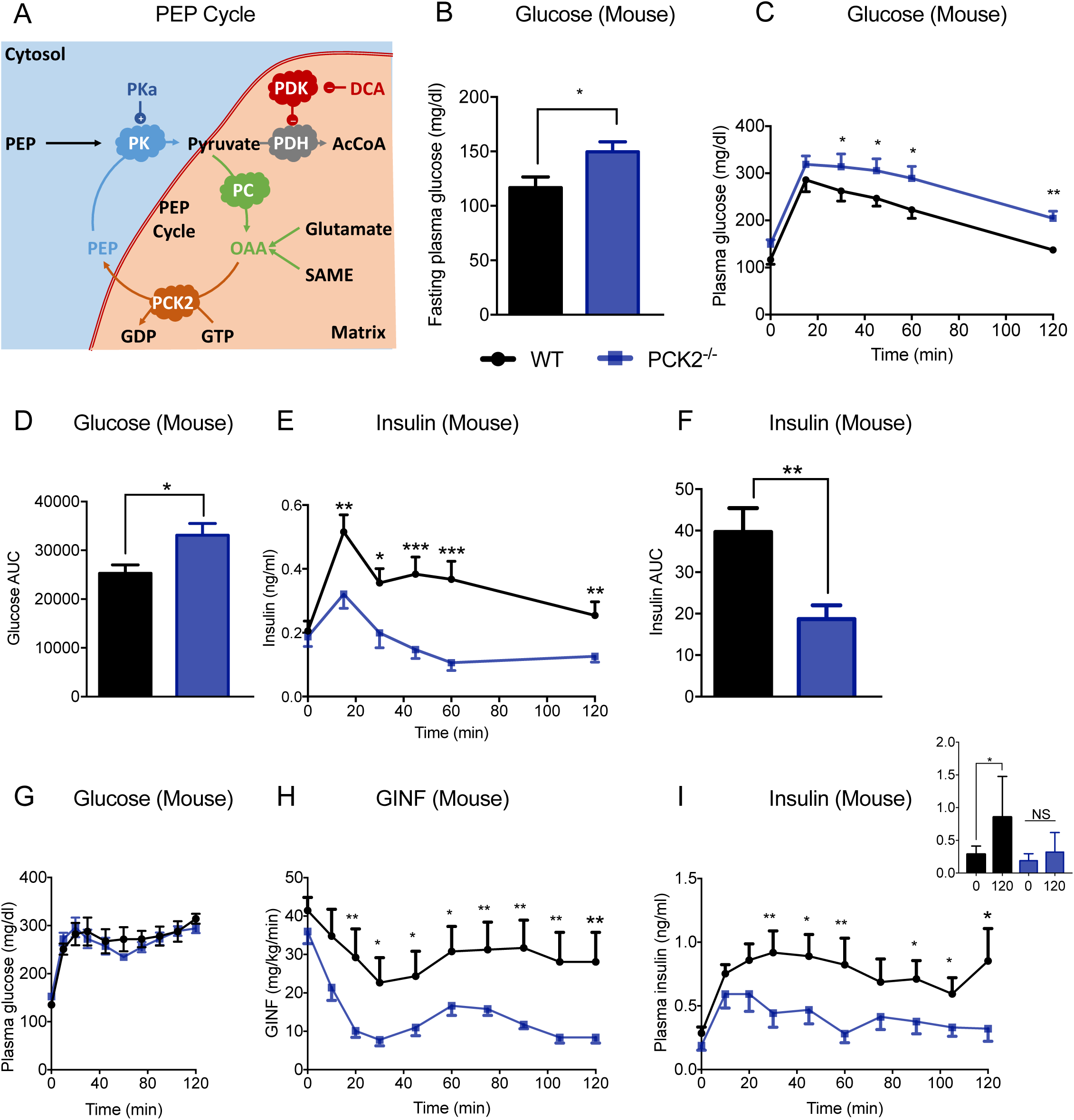
PCK2^-/-^ mice have impaired beta-cell function in vivo. **(A)** Schematic overview of strategies for assessing the OxPhos-independent mitochondrial GTP (mtGTP)-dependent phospho*enol*pyruvate (PEP) cycle. Oxidation of pyruvate by pyruvate dehydrogenase (PDH) is inhibited by phosphorylation by PDH Kinase (PDK, in **red**) that is in turn inhibited by dichloroacetic acid (DCA) or by knockout. Anaplerotic synthesis (**green arrows**) of oxaloacetic acid (OAA) is from pyruvate carboxylase (PC) or via metabolism of succinic acid methyl ester (SAME) or glutamate. OAA cataplerosis of OAA to generate PEP (**orange arrow**) is through the GTP-dependent PCK2 reaction. Cytosolic and mitochondrial PEP are hydrolyzed via pyruvate kinase (PK) that is allosterically enhanced with PK activators (**blue**). Glucose homeostasis in control (**black**) and PCK2^-/-^ (**blue**) mice after an (**B**) overnight fast, (**C-F**) during an OGTT, and (**G-I**) during a hyperglycemic clamp. (**B**) Basal, (**C**) stimulated, and (**D**) area under the curve (AUC) plasma glucose following an OGTT. (**E**) Simulated and (**F**) AUC insulin from the OGTT. (**G**) Plasma glucose concentrations, **(H)** Glucose Infusion Rate (GIR), and (**I**) insulin during the hyperglycemic portion of the clamp study. Mice were overnight fasted before the glucose tolerance test and hyperglycemic clamp. Data are represented as mean ± SEM (OGTT: n=11 per group; Hyperglycemic clamp: n=8 per group). Statistical comparisons (*, P <0.05; **, P<0.01; n.s., not significant) made by t-test.

## Results/Discussion

### Enhancing oxidative metabolism does not improve islet function

The final step of glycolysis generates pyruvate which, due to the restricted expression of lactate dehydrogenase (LDH) from beta-cells, only has two potential dispositions: entry into oxidative metabolism via pyruvate dehydrogenase (PDH), or entry into anaplerotic pathways via PC (Schuit et al., 2012). PDH kinase (PDK) phosphorylates PDH to inhibit PDH activity. Inhibiting PDK with the cell permeable dichoroacetate (DCA) reduces phosphorylation of PDH and, in principle, provides an acute pharmacological way to increase the oxidation of glucose (Sharma et al., 2019). Rather than improving glucose-stimulated insulin secretion (GSIS) by activating PDH, we confirmed prior *in vivo* and *in vitro* reports (Akhmedov et al., 2012) that DCA reduces insulin secretion in rat islets and rat insulinoma cells (INS-1) (**Figures S1A- S1C**). Mice with knockout of both isoform 2 and 4 of PDK (PDKdKO) mice have increased glucose oxidation rates as well as improved glucose tolerance, but paradoxically at the same time reduced plasma insulin levels (Wu et al., 2018). PDKdKO mice were utilized to evaluate specifically whether increasing the oxidative potential by reducing PDK activity had a similar effect to DCA in intact islets without the off-target effects of DCA (James et al., 2017; Jeoung et al., 2012). Perifused islets from PDKdKO mice had normal first phase insulin secretion, but the mitochondria-dependent second phase was significantly decreased (**Figures S1D and S1E**). Together, these DCA and PDKdKO data suggest that activating glucose OxPhos through inhibition of PDK would not be an effective therapeutic strategy to improve islet function.

### PCK2^-/-^ mice have impaired beta-cell function *in vivo*

Compared to oxidative flux through PDH, anaplerosis through pyruvate carboxylase (PC) is more steeply correlated with insulin secretion (Alves et al., 2015; Cline et al., 2004; Lu et al., 2002; MacDonald and Chang, 1985b; Newgard et al., 2002; Pongratz et al., 2007; Prentki et al., 2013). In principle, pyruvate oxidation, the cornerstone of the consensus model, does not change the concentration of TCA intermediates because OAA is regenerated with each turn of the cycle. Anaplerosis expands the TCA metabolite pool size through net synthesis of OAA while cataplerosis counterbalances this metabolite excess by exporting metabolites to shrink the TCA pool size. The mechanistic importance of pyruvate anaplerosis has been biochemically linked to the cataplerosis of OAA for the net synthesis of PEP in the PEP cycle (Jesinkey et al., 2019; Stark et al., 2009). Since beta-cells do not have cytosolic PEPCK (PCK1), non-glycolytic synthesis of PEP is dependent on the reaction of mtGTP with anaplerotic OAA via PCK2 (**Figure 1A**) (Jesinkey et al., 2019; MacDonald and Chang, 1985a; MacDonald et al., 1992; Stark et al., 2009). The importance of mtGTP synthesis was demonstrated *in vivo* and *in vitro* (Jesinkey et al., 2019; Kibbey et al., 2007), still the involvement of cataplerosis via PCK2 in beta-cell function has only been supported in insulinoma cells *in vitro* (Jesinkey et al., 2019; Stark et al., 2009). Whole body PCK2^-/-^ mice were generated and backcrossed onto the C57Bl6j background. Knockout was confirmed by tissue mRNA, as well as islet protein and enzyme activity (**Figures S1F-S1H**). At 8 weeks of age the male mice have similar body weights as well as fasting plasma glucagon levels (**Figures S1I and S1J**). As a sign of disrupted glucose homeostasis, fasting plasma glucose was elevated even on a regular chow diet (**Figure 1B**). PCK2^-/-^ mice demonstrated glucose intolerance during an oral glucose tolerance test (OGTT) (**Figures 1C and 1D**) that could be explained, at least in part, by lack of an insulin secretion response (**Figures 1E and 1F**). To assess islet function more directly a hyperglycemic clamp was performed in the PCK2^*-/-*^ mice which required a higher glucose infusion rate (GIR) to maintain glycemia at 300 mg/dl (**Figures 1G and 1H**). The PCK2^*-/-*^ mice were deficient in first- and second-phase insulin secretion accounting for at least part of the increased GIR and consistent with an essential role of PCK2 in glucose sensing (**Figure 1I**).

### Both the PEP cycle and PK activation depend on PCK2

Islet intrinsic effects on beta-cell function from PCK2^*-/-*^ mice were assessed directly via islet perifusions. Islet insulin content was not significantly different from PCK2^*-/-*^ compared to controls (**Figure S1K)**, but GSIS from PCK2^*-/-*^ islets confirmed the *in vivo* findings of reduced secretion with first- and second-phase GSIS compared to littermate controls (**Figures 2A-D**) as well as KCl-depolarized, K_ATP_-channel independent secretion being lower (**Figures 2A and 2B**). In the accompanying manuscript, acute application of the PK activator 10 μM TEPP46 (PKa) to INS1 832/13 cells as well as mouse and human islets amplified insulin release (Lewandowski et al., 2019). Here, PKa only stimulated insulin secretion from the controls identifying a dependence on PCK2-generated-PEP for the PKa-mediated effect (**Figures 2C and 2D**).

**Figure 2.**
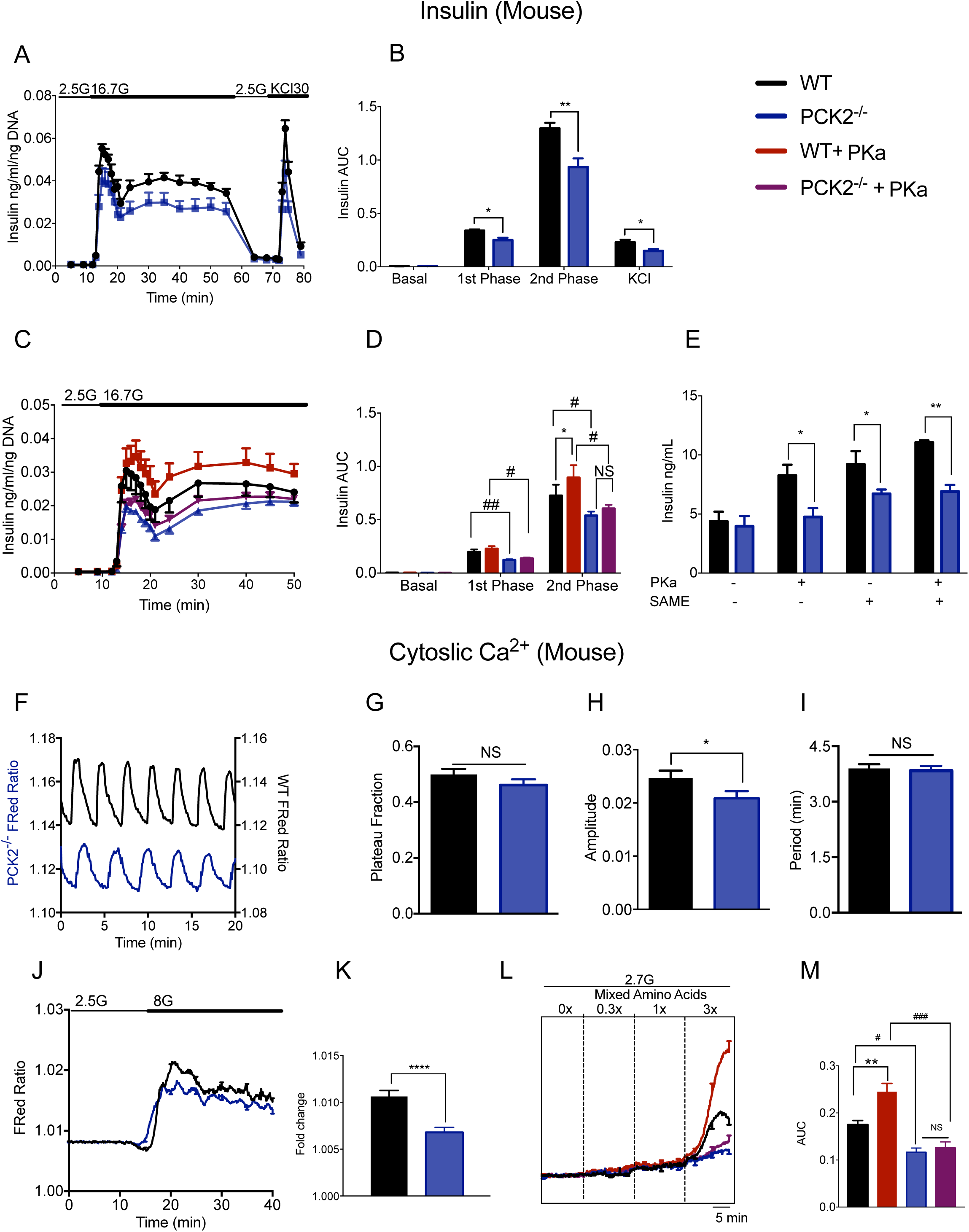
PCK2 is required for the anaplerosis and PK-dependent secretory response. **(A)** and (**B**) phasic insulin secretion during perifusion from 2.5→16.7 mM glucose of islets from control (WT) and PCK2^-/-^ (N=4). **(C)** and (**D**) phasic insulin secretion during perifusion from 2.5→16.7 mM glucose of islets from control (WT) and PCK2^-/-^ with or without treatment with 10 μM PKa (Tepp46) (N=4). **(E)** Insulin secretion from isolated, dispersed and reaggregated islets ± 10 μM PKa ± 10 mM SAME (N=4). **(F-M)** Cytosolic calcium recordings, in the presence of indicated glucose and amino acids concentrations from control (**black**) and PCK2^-/-^ (**blue**) islets. (**F**) Detrended calcium traces, (**G**) the fraction of time spent in the active phase (plateau fraction), and (**H**) average calcium amplitude from islets held at 8 mM glucose in physiologic amino acids (n=3 mice/group). **(I)** Period time between WT and PCK2-/- mice. **(J-K)** Ca2^+^ response following an acute increase glucose in islets from WT and PCK2^-/-^ mice. **(L-M)** Cytosolic Ca^2+^ in response to amino acids at 2.7 mM glucose ± 10 μM TEPP46 (PKa) (N=8). Data are represented as mean ± SEM. Statistical comparisons made by t-test and by 2-way ANOVA (*, P<0.05; **, P<0.01, ***, P<0.001, **#**, P<0.05, **##**, P<0.01).

### PCK2 is required for the anaplerotic and PKa-dependent insulin secretory response

The cell permeable methyl ester of succinate (SAME) is exclusively an anaplerotic substrate unlike glucose which supplies *both* oxidizing and anaplerotic pathways (Alves et al., 2015). SAME metabolism in beta-cells potently stimulates insulin secretion *in vitro* and *in vivo* (MacDonald and Fahien, 1988; Malaisse-Lagae et al., 1994; Zawalich et al., 1993). In control mouse islets, SAME-stimulated insulin release was further potentiated by the PK activator (**Figure 2E**) similar to that seen in human islets (Lewandowski et al., 2019). Just as with glucose, SAME did not stimulate as much insulin release from PCK2^*-/-*^ islets as controls and did not synergize with PK activation, an indication that anaplerosis is dependent upon PCK2 for the PK-dependent salutary response. Remarkably, despite poor GSIS, PCK2^-/-^ islets held at 8 mM glucose, akin to second-phase secretion, displayed a similar cytosolic calcium duty cycle and oscillation frequency to control islets, though the amplitude levels were slightly but significantly decreased (**Figures 2F-2I**). Either these small changes were sufficient to impair secretion, or this is also evidence of an additional role for PCK2 distal to cytosolic calcium. The calcium response following an acute increase in glucose is more indicative of first-phase secretion and was also diminished in PCK2^-/-^ relative to controls (**Figures 2J and 2K**). The calcium response in mouse islets at low glucose is especially sensitive to anaplerotic amino acids, a response that is muted as the glucose concentration dominates the response at higher concentrations (Lewandowski et al., 2019). Here, control islets had the expected strong calcium response to an acute increase in amino acids that was further enhanced by PKa (**Figures 2L and 2M**). PCK2^-/-^ islets did not respond either to amino acids or to PKa suggesting that both PK activation and anaplerosis require PCK2 to stimulate insulin secretion.

### PK activation improves insulin secretion in rodent and human models of T2DM

PK activation improved insulin secretion from insulinoma cells as well as mouse and rat islets and healthy human islets in a glucose-dependent manner (Lewandowski et al., 2019) To assess potential translatability as a beta-cell enhancer in disorders of glucose homeostasis, we interrogated whether PK activation might improve islet function in diabetes models. Insulin secretion was improved in a glucose-dependent manner from islets isolated from male rats fed a 3-week HFD and then acutely treated with a structurally distinct, cell permeable and orally bioavailable PK activator, AG519 (PKa2) (**Figure 3A**). PKa2-treated, KCl-depolarized islets further amplified insulin secretion (**Figure S2A**) indicative of the contribution of PK activation in K_ATP_-independent mechanisms. To determine the durability of the PK activator response, a second set of rats were fed HFD for three weeks with or without concomitant twice daily oral PKa2. Islets were isolated after an overnight fast without any additional PK activator treatment (either *in vivo* or *following islet isolation)*. Despite the absence of dosing for 16 hours, the chronically PKa2 treated islets displayed glucose-dependent amplification of insulin release **(Figure 3B)** similar to acute treatment without a change in islet insulin content **(Figure S2B)** suggesting either residual PKa2-mediated activation or *in vivo* protection from HFD-induced metabolic dysfunction.

**Figure 3.**
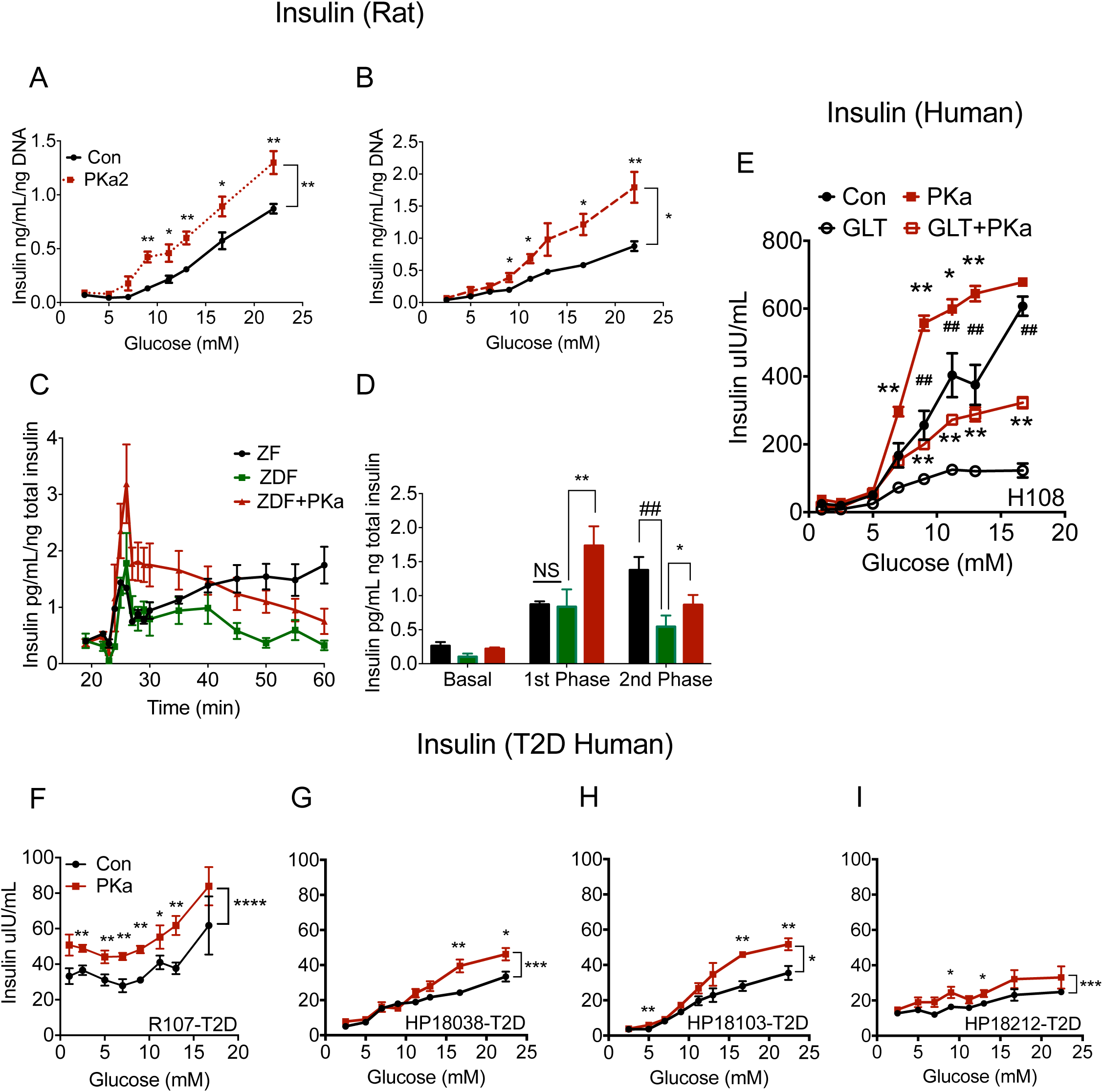
PK activation improves insulin secretion in rodent and human islet models of insulin resistance and T2DM. Insulin response to glucose concentration in islets from 3-week HFD rats (**A**) without (black) or with (red, dotted) acute *in vitro* 10 μM PKa2 (AG519) (N=4) and (**B**) with twice daily peanut butter vehicle without (black) or with 25 mg/kg PKa2 (N=4) but without exposure to drug for 16 hours. (**C**) and (**D**) phasic insulin secretion from perifused islets (2.5 mM→16.7 mM glucose) from regular chow (ZF, **black**) and high fat fed (ZDF) Zucker rats without (**green**) and with (**red**) 10 μM PKa (N=4). **(E)** Insulin response to glucose concentration from healthy human reaggregated islets (donor H108) treated for 72 h with BSA control (**closed**) vs. glucolipotoxic (GLT, 20 mM glucose and 1 mM 2:1 oleate:palmitate) conditions (**open**) conditions without (**black circles**) or with (**red squares**) 10 μM PKa). **(F-I)** Insulin response to glucose concentration in reaggregated T2D human (see Table S1for characteristics) islets without (**black**) or with (**red**) 10 μM PKa. Data are represented as mean ± SEM. Statistical comparisons made by t-test and by 2-way ANOVA (*with vs. without PKa; ^#^, ZF vs. ZDF or control vs. GLT; *,^#^, P<0.05; **,^##^, P<0.01).

Although a HFD increases insulin resistance and causes dysglycemia, it does not precipitate diabetes in normal rats. Therefore, we used the Zucker rat which is deficient in the leptin receptor and develops diabetes on a HFD (ZDF) compared to lean controls (ZF) **(Figure S2C)**. ZDF islets had lower islet insulin content compared to ZF controls **(Figure S2C)** and had markedly diminished second-phase insulin secretion during the glucose perifusion (**Figures 3C and 3D**). Notably, acute treatment of diabetic ZDF islets with PKa improved both first- and second-phase insulin secretion (**Figure 3C and 3D**).

As an *in vitro* model of diabetic dysglycemia, normal human islets incubated for 72-hours under glucolipotoxic (GLT) conditions (20 mM glucose + 1 mM 2:1 oleate:palmitate) had diminished GSIS compared to normal culture conditions (**Figure 3E**). Acute treatment of control human islets with PK activator augmented insulin secretion in a glucose-dependent manner as in Lewandowski et al. (Lewandowski et al., 2019). Treatment of GLT-exposed islets with PKa improved glucose-dependent insulin secretion to the level of the untreated control islets. As a more direct assessment of the potential translatability of PK activation to humans with T2DM, PK improved GSIS islets from 4 different *bona fide* T2D donors **(Figures 3F-3I).** In all of these diabetes models, PK enzymatic activity in islet lysates was not deficient and was fully activatable by PK activator **(Figures S2D-S2F).** As was seen in insulinoma cells (Lewandowski et al., 2019), multiple chemically distinct PK activators amplified insulin release from human islets **(Figure S2G)**, reducing the likelihood of a single off-target mechanism to explain the improvement in islet function. Taken together, these *in vitro* data in multiple diabetes models using two chemically distinct PK activators, suggest PK activation may be a targetable mechanism to improve islet function and potentially beta-cell health.

### PK activation improves insulin secretion and health *in vivo*

The glucose-dependency of PK activation was assessed *in vivo*. Following an overnight fast, healthy male rats were given a primed, continuous infusion of vehicle or either PKa or PKa2 with target plasma concentration of ∼10 μM (**Figure S3A**). Neither fasting glucose or insulin were altered in response to the drug infusion (**Figure 4A**). Following 90 minutes of continuous drug exposure, glucose concentrations were gently ramped to a peak of 250 mg/dl by a variable glucose infusion over 70 minutes (**Figure 4A**). Both PK activators significantly improved insulin secretion once glucose was raised above 150 mg/dl (**Figures 4B and 4C**). A linear regression of all the individual glucose and insulin samples from the rats showed a doubling of GSIS at all points above this threshold (**Figure S3B**). Consistent with higher insulin secretion, glucose infusion rate (GINF) was higher in PK activator treated rats (**Figure S3C**). HFD-fed rats were treated for three weeks with twice daily oral vehicle or PKa2 and then subjected to an overnight fast in the absence of further drug administration. A hyperglycemic ramp was performed over 50 minutes. Fasting and ramped plasma glucose concentrations were identical between both groups (**Figure 4D**). Glucose infusion rate (GINF) was not changed in HFD-fed rats that were treated for three weeks with twice daily oral vehicle or PKa2 (**Figure S3E).** In contrast, prior PKa2 treatment nearly doubled fasting and stimulated insulin levels compared to controls (**Figure 4E and 4F**) and was independent of bodyweight and plasma glucagon (**Figures S3D and S3F**). Similar to the islets from the three-week HFD fed rats (**Figure 3B**), the improved function may be a consequence of improved islet health and/or residual PK activation in the *absence* of treatment for 24-hours. Taken together, PCK2-dependent activation of the PEP cycle via two chemically distinct orally available small molecules improved islet function in preclinical diabetes models, in diabetic human islets and up to three weeks *in vivo*.

**Figure 4.**
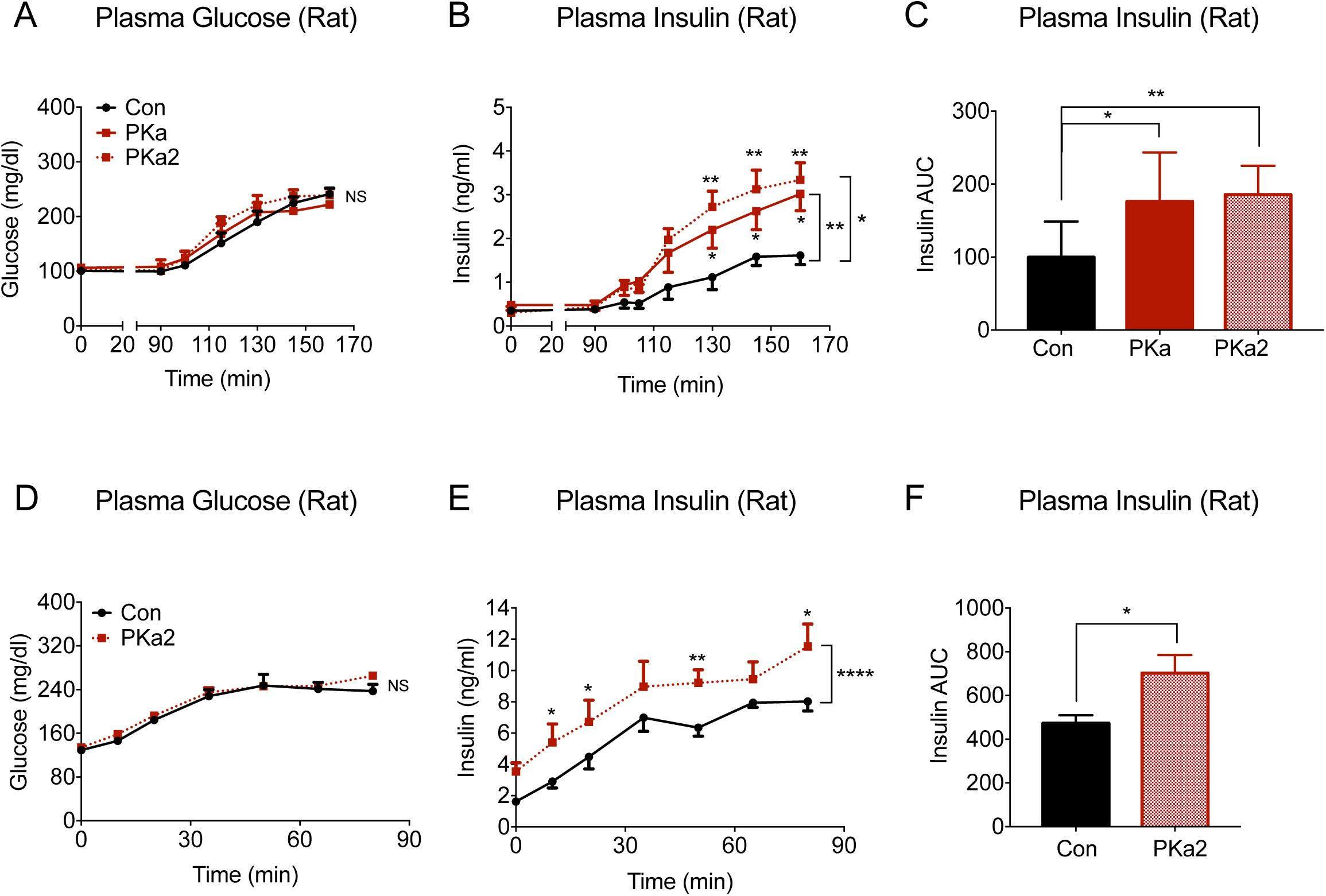
PK activation improves GSIS in regular chow and HFD-fed rats. (**A**) Plasma glucose, (**B**) plasma insulin, and (**C**) insulin AUC after primed-continuous intravenous infusion (t=0) followed by hyperglycemic ramp (t = 90-160 min) of PKa1, PKa2, vs. vehicle in over-night fasted regular chow fed rats (N=6-7 per group). (**D**) Plasma glucose, (**E**) plasma insulin, and (**F**) insulin AUC in 3-week HFD rats with twice daily administration of peanut butter vehicle without (black) or with PKa2 (red) then subjected to a hyperglycemic ramp after an overnight fast without subsequent drug treatment (n= 6-7 per group). Statistical comparisons made t-test and by 2-way ANOVA (*, P<0.05, **, P<0.01).

## Conclusions

PK activation by small molecules improves insulin secretion in normal and diabetes models from insulinomas, mouse, rat, and human islets in a glucose-dependent manner. This glucose-dependency minimizes the risk of hypoglycemia similar to GLP1R agonists but distinct from the increased risk from GK activation or sulfonylurea receptor agonism. A benefit of PK activation would not have been predicted by the consensus mechanism of beta-cell glucose sensing, where GK is proposed to determine the extent of OxPhos. Instead, this PK-dependent glucose-sensing is mechanistically dependent on anaplerotic metabolism (e.g., pyruvate carboxylase, amino acids, or SAME) coupled to PK through PEP produced by the mitochondrial GTP-dependent enzyme PCK2. As shown in the companion manuscript (Lewandowski et al., 2019), the PK catalyzed reaction harnesses the free energy of PEP hydrolysis to cause ADP privation of both the mitochondria and K_ATP_ channels to depolarize the plasma membrane. The PK activator AG348 has been demonstrated to be well tolerated in Phase 1 healthy volunteer studies (Yang et al, 2018) and a Phase 2 trial in patients with PK deficiency (Grace et al., 2019; Yang et al., 2019). Our work does not identify an underlying PK deficiency *per se* contributing to islet dysfunction in T2D and may even argue against it. Nevertheless, these data provide supporting preclinical data that PK activation or other mechanisms that accelerate the PEP cycle may be beneficial targets in diseases such as T2DM where glucose metabolism is aberrant.

## ACKNOWLEDGEMENTS

We thank Dr. Robert A. Harris and Yasmeen Rahimi for providing mice lacking pyruvate dehydrogenase kinase 2 and 4 (PDK double knock out; PDKDKO). The Kibbey laboratory gratefully acknowledges NIH/NIDDK (R01DK092606, R01DK110181 and K08DK080142), CTSA (UL1RR-0024139), and DRC (P30DK045735) as well as an investigator-initiated Sponsored Research Agreement from Agios Pharmaceuticals to Yale supporting this work. This work was supported in part by a Monash-Yale Strategic Grant Program in Metabolism to RS and RGK (MDO17PW01). Romana Stark was supported by a National Health and Medical Research Council of Australia Early Career Fellowship. Zane B Andrews was supported by a National Health and Medical Research Council of Australia Senior Research Fellowship (APP1154974). The Merrins laboratory gratefully acknowledges support from the American Diabetes Association (1-16-IBS-212), the NIH/NIDDK (K01DK101683 and R01DK113103), the NIH/NIA (R21AG050135 and R01AG062328), the Wisconsin Partnership Program, and the Central Society for Clinical and Translational Research.

## AUTHOR CONTRIBUTIONS

RGK conceived the study and wrote the paper with AA, RS and RLC. AA and RLC performed the main body of experiments with contributions from RS, TCA, and SLL and assisted by BW, XZ and SS. CK, CLT, RS, ZBA provided reagents and technical expertise. RGK, MJM, RS, ZBA obtained funding. RGK, AA, RLC, RS, SLL, MJM, CLT, and CK interpreted the data and edited the manuscript.

## Materials and Methods

### Experimental model and subject details

All animal studies were approved by the Yale University Institutional Animal Care and Use Committee and were performed in accordance with all regulatory standards. 240-260g male Sprague-Dawley rats were obtained from Charles River Laboratories (Wilmington, MA) and were single housed while they were fed either a high fat diet (Dyets #112245, Bethlehem, PA; 59% calories from fat, 26% from carbohydrate, 15% from protein) ad lib for 3 weeks or a chow diet (Harlan Teklad #2018, Madison, WI; 18% calories from fat, 58% from carbohydrate, 24% from protein). High fat diet (HFD) fed rats were treated twice daily for 3 weeks with PK activators (25 mg/kg) in peanut butter or peanut butter vehicle while they were maintained on the HFD diet. Detailed information of PKa activators, PKa (TEPP46) and PKa2 (AG519), are provided in the key resource table. Rats underwent the placement of jugular venous catheters for blood sampling and carotid artery for infusion ∼7 days before the terminal studies and recovered their pre-surgical weights by 5-7 days after the operation. All infusions were performed on overnight fast rats. At the end of each study, rats were euthanized by IV pentobarbital. Mice received ad libitum access to food and water and were bred, maintained and housed in the Yale Animal Resources Center at 23°C under 12-hour light/12-hour dark cycles (0700–1900h). All strains were on a C57BL/6J background and male mice were studied at 8 to 9 weeks of age. All of the mice studied were fed on a regular chow diet (Harlan Teklad TD2018: 18% fat, 58% carbohydrate, and 24% protein). Study cohorts consisted of homozygous PCK2^-/-^ mice and littermate male WT controls. Catheters were placed in the jugular vein 6–9 days before the experiments; only mice that recovered more than 95% of their pre-operative body weight were studied.

### PCK2^-/-^ mouse

Heterozygous PCK2 knockout mice (denoted C57BL/6NTac-Pck2^tm1 (KOMP)Vlcg^, project ID VG13655) were obtained from the University of California Davis Knockout Mouse Project repository. For these knockout mice the VelociGene targeting strategy was employed, which enables the rapid and high-throughput generation of custom gene mutations in mice according to method described previously (Dechiara et al., 2009; Valenzuela et al., 2003). The insertion of Velocigene cassette ZEN-Ub1 created a deletion of size 8372bp between positions 55548778-55540407 of Chromosome 14. Mice harbor a gene expression reporter (*lacZ*), and mice had a neomycin cassette. At the Monash Animal Research Facility heterozygous knockout mice were crossed with CMV-Cre mice, which express Cre recombinase under the CMV promoter, to delete the loxP flanked selection cassette, and to generate global neomycin cassette-free PCK2 knockout mice. Heterozygous PCK2 knockout mice were further crossed with C57BL6J mice to create PCK2 knockout mice on a C57BL6J background. Offspring that inherited the targeted PCK2 allele were interbred as heterozygotes for production of experimental PCK2^+/+^ and PCK2^-/-^ mice. PCK2^-/-^ mice behave normally and are viable and fertile.

### Hyperglycemic- clamp studies

In hyperglycemic clamp studies, overnight-fasted, awake mice under gentle tail restraint were infused with 20% D- glucose at a variable rate to maintain hyperglycemia (240–260 mg/dl). Plasma glucose was measured using a Analox GL5 Analyzer (Analox Instruments Ltd., UK). Serum insulin was measured via Mouse Ultrasensitive ELISA (80-INSMSU-E01, ALPCO). Fasting serum glucose and insulin concentration were also measured using the same methods, respectively.

#### For acute PKa treatment and hyperglycemic clamp

One week prior to experimentation, chow-fed male SD rats (∼300g) underwent surgical arterial and venous catheterization. After recovering from surgery, rats were starved overnight and received an intravenous bolus of compound PKa (11mg/kg) or vehicle control followed by a continuous infusion with compound PKa 0.17umol/kg/min throughout the entire infusion study. At the point of 90 minutes, 20% dextrose was infused at a variable rate to reach hyperglycemia (240–260 mg/dl). Blood samples (50 μl) were taken every 15 min for measurement of glucose using a YSI glucose analyzer.

#### For chronic PKa treatment and hyperglycemic clamp

HFD-fed rats treated with PKa were overnight fasted to start the experiment and the PKa in peanut butter was given before the overnight fasting. Plasma samples were collected before the start of infusion (basal 0 minute) to measure glucose and insulin levels. Rates of infused 20% glucose was variably increased as shown to reach target hyperglycemia while collecting plasma samples at 15, 30, 45, 60, 75, 90 minutes.

### Oral glucose tolerance test (OGTT)

Glucose tolerance tests were performed after an overnight fast. Mice were gavaged with 1 g/kg glucose, and blood was collected by tail bleed at 0, 15, 30, 45, 60, 90, and 120 min for plasma insulin and glucose measurements. Plasma glucose concentrations were measured using the glucose oxidase method on an Analox GL5 Analyzer (Analox Instruments Ltd., UK). Serum insulin was measured via Mouse Ultrasensitive ELISA (80-INSMSU-E01, ALPCO). Fasting serum glucose and insulin concentration were also measured using the same methods, respectively.

### Islet isolation and GSIS

Pancreata were excised from anesthetized rodents (rats and mice) and islets were isolated by collagenase P digestion followed by centrifugation with Histopaque 1100 solution for density separation of pure islets and then hand-picked as previously described (Kibbey et al., 2014). Islets were allowed to recover for 24-48 hours in RPMI with 10% FBS and antibiotics with either 5mM glucose (rats) or 11mM glucose (mice). Approximately 50-80 islets were then layered between a slurry of acrylamide gel column beads (Bio-Gel P4G; BioRad150-412) and perifusion buffer DMEM (D-5030, Sigma) supplemented with 24mM NaHCO_3_, 2.5 mM glucose, 10 mM HEPES, 2 mM glutamine and 0.2% fatty acid free BSA. The islets were perifused at 100 ul/min for a 1hr equilibration period using a Bio-Rep Perifusion Instrument (Miami, FL) that maintains precise temperature, gas (5% CO2/95% air) and flow control. After the stabilization period, the islets were perifused with 2.5 mM glucose for 10 minutes followed by stimulatory 16.7 mM glucose for 45 minutes followed by 30 mM KCl as indicated. During the perifusion, outflow was collected into a 96-well plate format and secreted insulin concentrations were measured by a high range rodent insulin ELISA assay kit (ALPCO) and normalized to islet DNA which was measured using the Quant-iT PicoGreen dsDNA Reagent (Life Technologies Corporation).

### Timelapse Imaging

For measurements of cytosolic Ca^2+^, islets were pre-incubated in 2.5 µM FuraRed (Molecular Probes F3020) in islet media for 45 min at 37°C before they were placed in a glass-bottomed imaging chamber (Warner Instruments) mounted on a Nikon Ti-Eclipse inverted microscope equipped with a 20X/0.75NA SuperFluor objective (Nikon Instruments). The chamber was perfused with a standard external solution containing 135 mM NaCl, 4.8 mM KCl, 2.5 mM CaCl_2_, 1.2 mM MgCl_2_, 20 mM HEPES (pH 7.35). The flow rate was 0.4 mL/min and temperature was maintained at 33°C using solution and chamber heaters (Warner Instruments). Excitation was provided by a SOLA SEII 365 (Lumencor) set to 10% output. Single DiR images utilized a Chroma Cy7 cube (710/75x, T760lpxr, 810/90m). Excitation (430/20x and 500/20x) and emission (630/70m) filters (ET type; Chroma Technology) were used in combination with an FF444/521/608-Di01 dichroic beamsplitter (Semrock) and reported as the excitation ratio (R430/500). Fluorescence emission was collected with a Hamamatsu ORCA-Flash4.0 V2 Digital CMOS camera every 6 s. A single region of interest was used to quantify the average response of each islet using Nikon Elements and custom MATLAB software (MathWorks).

### Human Islet Insulin Secretion Studies

Human islets from normal and Type 2 diabetic donors obtained from the University of Alberta Diabetes Institute, the University of Chicago Diabetes Research and Training Center, or Prodo Laboratories Inc. (CA) were cultured in glutamine-free CMRL supplemented with 10 mM niacinamide and 16.7 μM zinc sulfate (Sigma), 1% ITS supplement (Corning), 5 mM sodium pyruvate, 1% Glutamax, 25 mM HEPES (American Bio), 10% HI FBS and antibiotics (10,000 units/ml penicillin and 10 mg/ml streptomycin). All media components were obtained from Invitrogen unless otherwise indicated. Human donor islets were cultured intact, then dispersed and re-aggregated as pseudo-islets for dynamic insulin secretion studies. Islets were dispersed with Accutase (Invitrogen) and the resulting cell suspension was seeded at 5000 cells per well of a 96-well V-bottom plate, lightly centrifuged (200*g*) and then incubated for 12-24 h at 37°C 5% CO_2_/95% air for the islet to reform into an intact islet. Dynamic GSIS studies were performed 24 h after the islets were plated into the 96 well plates following dispersion and re-aggregation. The human islet plates were washed and incubated at 37°C 5% CO_2_/95% air in DMEM (Sigma D5030) supplemented with 24mM NaHCO_3_, 2.5mM glucose, 10 mM HEPES, 2 mM glutamine and 0.2% fatty acid-free BSA for 1.5 h. After the first incubation, human islet plates were then washed with glucose free study DMEM and incubated for 2 hours in DMEM study media with 1, 2.5, 5, 7, 9, 11.2, 13, 16.7 or 22.4 mM glucose in presence of 10 µM TEPP-46 or 0.1% DMSO. To induce glucolipotoxicity in normal donors, islet re-aggregates were incubated for 72 h under control or glucolipotoxic conditions (1% fatty acid free BSA alone or conjugated with 1mM of 2:1 sodium oleate/sodium palmitate and 20 mM glucose), which were removed and replaced with DMEM as described above for the acute study. Supernatants were evaluated by insulin ELISA (ALPCO).

## KEY RESOURCES TABLE

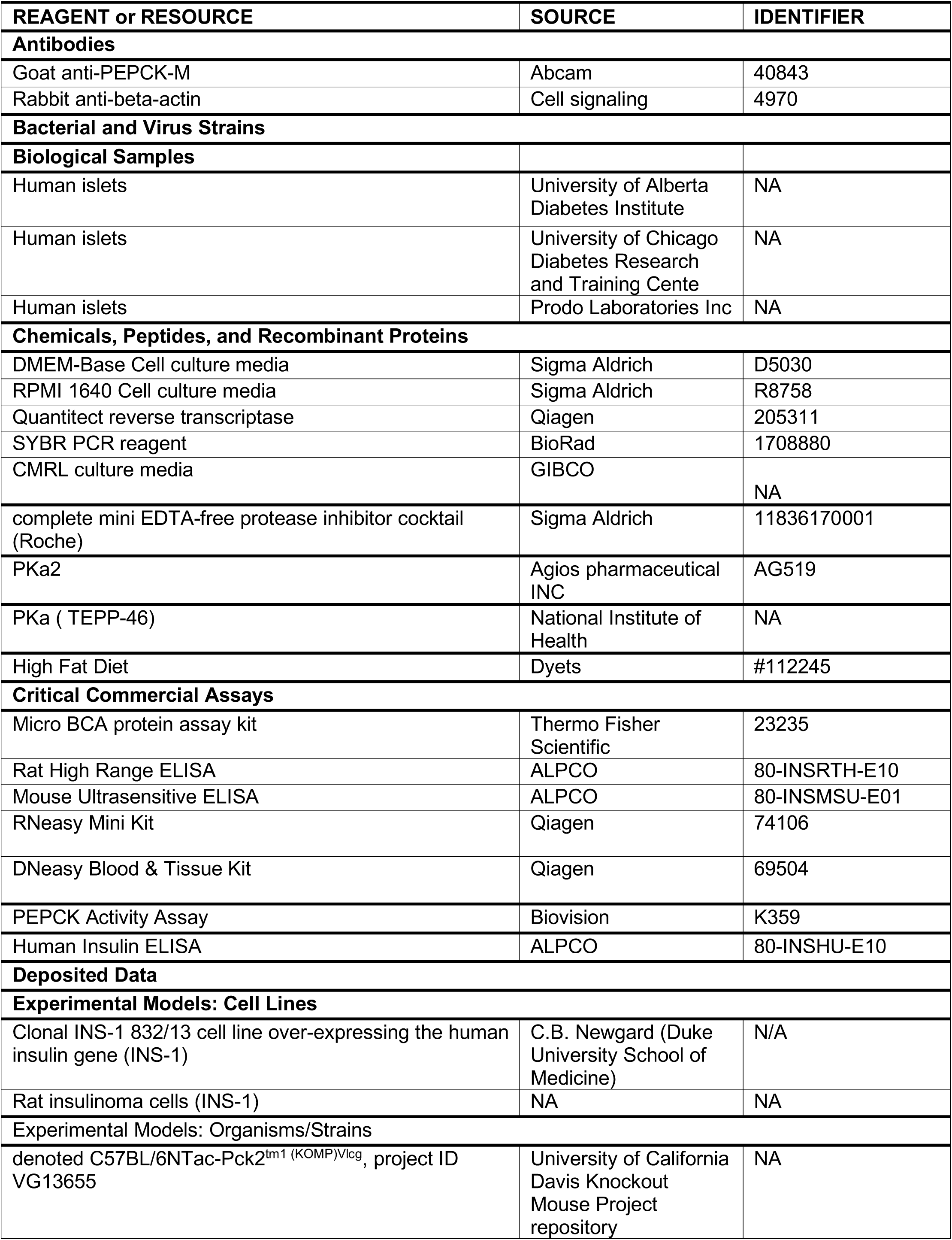
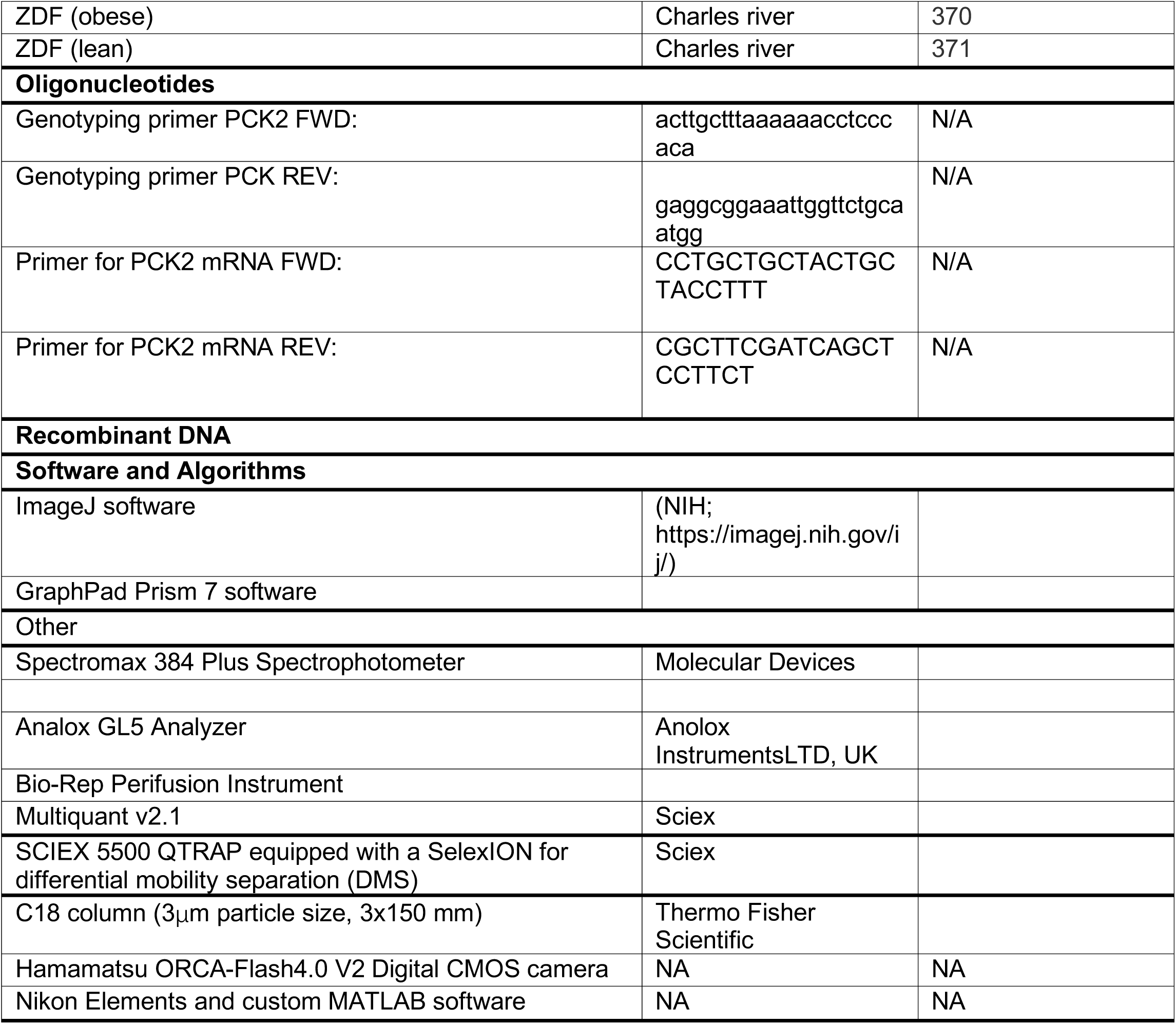

## Reference

Akhmedov, D., De Marchi, U., Wollheim, C.B., and Wiederkehr, A. (2012). Pyruvate dehydrogenase E1alpha phosphorylation is induced by glucose but does not control metabolism-secretion coupling in INS-1E clonal beta-cells. Biochim Biophys Acta 1823, 1815–1824.

Alves, T.C., Pongratz, R.L., Zhao, X., Yarborough, O., Sereda, S., Shirihai, O., Cline, G.W., Mason, G., and Kibbey, R.G. (2015). Integrated, Step-Wise, Mass-Isotopomeric Flux Analysis of the TCA Cycle. Cell Metab 22, 936–947.

Cline, G.W., Lepine, R.L., Papas, K.K., Kibbey, R.G., and Shulman, G.I. (2004). 13C NMR isotopomer analysis of anaplerotic pathways in INS-1 cells. J Biol Chem 279, 44370–44375.

De Ceuninck, F., Kargar, C., Ilic, C., Caliez, A., Rolin, J.O., Umbdenstock, T., Vinson, C., Combettes, M., de Fanti, B., Harley, E., et al. (2013). Small molecule glucokinase activators disturb lipid homeostasis and induce fatty liver in rodents: a warning for therapeutic applications in humans. Br J Pharmacol 168, 339–353.

Dechiara, T.M., Poueymirou, W.T., Auerbach, W., Frendewey, D., Yancopoulos, G.D., and Valenzuela, D.M. (2009). VelociMouse: fully ES cell-derived F0-generation mice obtained from the injection of ES cells into eight-cell-stage embryos. Methods Mol Biol 530, 311–324.

Efanova, I.B., Zaitsev, S.V., Zhivotovsky, B., Kohler, M., Efendic, S., Orrenius, S., and Berggren, P.O. (1998). Glucose and tolbutamide induce apoptosis in pancreatic beta-cells. A process dependent on intracellular Ca2+ concentration. J Biol Chem 273, 33501–33507.

Erion, D.M., Lapworth, A., Amor, P.A., Bai, G., Vera, N.B., Clark, R.W., Yan, Q., Zhu, Y., Ross, T.T., Purkal, J., et al. (2014). The hepatoselective glucokinase activator PF-04991532 ameliorates hyperglycemia without causing hepatic steatosis in diabetic rats. PLoS One 9, e97139.

Grace, R.F., Rose, C., Layton, D.M., Galacteros, F., Barcellini, W., Morton, D.H., van Beers, E.J., Yaish, H., Ravindranath, Y., Kuo, K.H.M., et al. (2019). Safety and Efficacy of Mitapivat in Pyruvate Kinase Deficiency. N Engl J Med 381, 933–944.

Iwakura, T., Fujimoto, S., Kagimoto, S., Inada, A., Kubota, A., Someya, Y., Ihara, Y., Yamada, Y., and Seino, Y. (2000). Sustained enhancement of Ca(2+) influx by glibenclamide induces apoptosis in RINm5F cells. Biochem Biophys Res Commun 271, 422–428.

James, M.O., Jahn, S.C., Zhong, G., Smeltz, M.G., Hu, Z., and Stacpoole, P.W. (2017). Therapeutic applications of dichloroacetate and the role of glutathione transferase zeta-1. Pharmacol Ther 170, 166–180.

Jeoung, N.H., Rahimi, Y., Wu, P., Lee, W.N., and Harris, R.A. (2012). Fasting induces ketoacidosis and hypothermia in PDHK2/PDHK4-double-knockout mice. Biochem J 443, 829–839.

Jesinkey, S.R., Madiraju, A.K., Alves, T.C., Yarborough, O.H., Cardone, R.L., Zhao, X., Parsaei, Y., Nasiri, A.R., Butrico, G., Liu, X., et al. (2019). Mitochondrial GTP Links Nutrient Sensing to beta Cell Health, Mitochondrial Morphology, and Insulin Secretion Independent of OxPhos. Cell Rep 28, 759–772 e710.

Kibbey, R.G., Choi, C.S., Lee, H.Y., Cabrera, O., Pongratz, R.L., Zhao, X., Birkenfeld, A.L., Li, C., Berggren, P.O., Stanley, C., et al. (2014). Mitochondrial GTP insensitivity contributes to hypoglycemia in hyperinsulinemia hyperammonemia by inhibiting glucagon release. Diabetes 63, 4218–4229.

Kibbey, R.G., Pongratz, R.L., Romanelli, A.J., Wollheim, C.B., Cline, G.W., and Shulman, G.I. (2007). Mitochondrial GTP regulates glucose-stimulated insulin secretion. Cell Metab 5, 253–264.

Lewandowski, S.L., Cardone, R.L., Foster, H.R., Ho, T., Evgeniy, P. C., VanDeusen, H.R., Alves, T.C., Zhao, X., Capozzi, M.E., Jahan, I., et al. (2019). Pyruvate kinase controls signal strength in the insulin secretory pathway. Cell metabolism

Lu, D., Mulder, H., Zhao, P., Burgess, S.C., Jensen, M.V., Kamzolova, S., Newgard, C.B., and Sherry, A.D. (2002). 13C NMR isotopomer analysis reveals a connection between pyruvate cycling and glucose-stimulated insulin secretion (GSIS). Proc Natl Acad Sci U S A 99, 2708–2713.

MacDonald, M.J., and Chang, C.M. (1985a). Do pancreatic islets contain significant amounts of phosphoenolpyruvate carboxykinase or ferroactivator activity? Diabetes 34, 246–250.

MacDonald, M.J., and Chang, C.M. (1985b). Pancreatic islets contain the M2 isoenzyme of pyruvate kinase. Its phosphorylation has no effect on enzyme activity. Mol Cell Biochem 68, 115–120.

MacDonald, M.J., and Fahien, L.A. (1988). Glyceraldehyde phosphate and methyl esters of succinic acid. Two “new” potent insulin secretagogues. Diabetes 37, 997–999.

MacDonald, M.J., McKenzie, D.I., Walker, T.M., and Kaysen, J.H. (1992). Lack of glyconeogenesis in pancreatic islets: expression of gluconeogenic enzyme genes in islets. Horm Metab Res 24, 158–160.

Malaisse-Lagae, F., Bakkali Nadi, A., and Malaisse, W.J. (1994). Insulinotropic response to enterally administered succinic and glutamic acid methyl esters. Arch Int Pharmacodyn Ther 328, 235–242.

Newgard, C.B., Lu, D., Jensen, M.V., Schissler, J., Boucher, A., Burgess, S., and Sherry, A.D. (2002). Stimulus/secretion coupling factors in glucose-stimulated insulin secretion: insights gained from a multidisciplinary approach. Diabetes 51 Suppl 3, S389–393.

Pongratz, R.L., Kibbey, R.G., Shulman, G.I., and Cline, G.W. (2007). Cytosolic and mitochondrial malic enzyme isoforms differentially control insulin secretion. J Biol Chem 282, 200–207.

Prentki, M., Matschinsky, F.M., and Madiraju, S.R. (2013). Metabolic signaling in fuel-induced insulin secretion. Cell Metab 18, 162–185.

Schuit, F., Van Lommel, L., Granvik, M., Goyvaerts, L., de Faudeur, G., Schraenen, A., and Lemaire, K. (2012). beta-cell-specific gene repression: a mechanism to protect against inappropriate or maladjusted insulin secretion? Diabetes 61, 969–975.

Sharma, G., Wu, C.Y., Wynn, R.M., Gui, W., Malloy, C.R., Sherry, A.D., Chuang, D.T., and Khemtong, C. (2019). Real-time hyperpolarized (13)C magnetic resonance detects increased pyruvate oxidation in pyruvate dehydrogenase kinase 2/4-double knockout mouse livers. Sci Rep 9, 16480.

Stark, R., Pasquel, F., Turcu, A., Pongratz, R.L., Roden, M., Cline, G.W., Shulman, G.I., and Kibbey, R.G. (2009). Phosphoenolpyruvate cycling via mitochondrial phosphoenolpyruvate carboxykinase links anaplerosis and mitochondrial GTP with insulin secretion. J Biol Chem 284, 26578–26590.

Valenzuela, D.M., Murphy, A.J., Frendewey, D., Gale, N.W., Economides, A.N., Auerbach, W., Poueymirou, W.T., Adams, N.C., Rojas, J., Yasenchak, J., et al. (2003). High-throughput engineering of the mouse genome coupled with high-resolution expression analysis. Nat Biotechnol 21, 652–659.

Wu, C.Y., Tso, S.C., Chuang, J.L., Gui, W.J., Lou, M., Sharma, G., Khemtong, C., Qi, X., Wynn, R.M., and Chuang, D.T. (2018). Targeting hepatic pyruvate dehydrogenase kinases restores insulin signaling and mitigates ChREBP-mediated lipogenesis in diet-induced obese mice. Mol Metab 12, 12–24.

Yang, H., Merica, E., Chen, Y., Cohen, M., Goldwater, R., Kosinski, P.A., Kung, C., Yuan, Z.J., Silverman, L., Goldwasser, M., et al. (2019). Phase 1 Single- and Multiple-Ascending-Dose Randomized Studies of the Safety, Pharmacokinetics, and Pharmacodynamics of AG-348, a First-in-Class Allosteric Activator of Pyruvate Kinase R, in Healthy Volunteers. Clin Pharmacol Drug Dev 8, 246–259.

Zawalich, W.S., Zawalich, K.C., Cline, G., Shulman, G., and Rasmussen, H. (1993). Comparative effects of monomethylsuccinate and glucose on insulin secretion from perifused rat islets. Diabetes 42, 843–850.

