## Supplemental data for "Pharmacologic activation of the mitochondrial phospho*enol*pyruvate cycle enhances islet function in vivo"

### Supplemental Information

#### Supplemental Figures

Figure S1 related to Introduction & Figure 1

Figure S2 related to Figure 3

Figure S3 related to Figure 4

#### Supplemental Tables

Table S1. Human islets donor characteristics related to figure 3.

Table S2. Physiological amino acid mixture concentrations, Related to Figure 2

#### SUPPLEMENTAL METHODS AND MATERIALS

**PEPCK activity assay:** PEPCK activity was measured according the protocol (Catalog # K359 Biovision). Simply described, 70-100 islets were homogenized with 200 μl cold PEPCK assay buffer provided in the kit for 10 minutes on ice, then centrifuged at 10,000× g at 4 °C for 10 minutes. Supernatants were then collected and protein concentration was measured. 50 ul of sample and 50μl of reaction mix consisting of PEPCK assay buffer, PEPCK converter, PEPCK developer, PEPCK probe and PEPCK substrate mix (Catalog # K359 Biovision) were added to each well and the plate was read at an OD of 570 nm.

**Immunoblotting analysis:** 200-250 islets were homogenized in ice-cold homogenization buffer with protease and phosphatase inhibitors (cOmplete MINI + PhosSTOP (Roche)). Proteins were detected by Western blot analysis after separating 30–50 µg of total protein lysate on a 4–12% Tris-glycine gel (BioRad) and transferring to a polyvinylidene difluoride membrane (Immobilon-P 0.45 µm, Millipore). Goat anti-PEPCK-M (Abcam) and rabbit anti-B-actin (cell signaling) were used.

**Quantitative PCR:** Total RNA was isolated from islets using buffer RLT and was further purified with the RNeasy kit (Qiagen). The abundance of transcripts was assessed by real-time PCR on an Applied Biosystems 7500 Fast Real-Time PCR System with a SYBR Green detection system (Bio-Rad). The PCK2 primer sequences are provided in key resource table.

**PK Activity Assay:** The enzymatic assay of pyruvate kinase (EC 2.7.1.40) protocol from Sigma was adapted to a 96 well plate format, and modified using a 1:2 PEP dilution curve starting at 2.5 mM or 5 mM PEP. Unless specified, 3 µM FBP was present.

**PDKdKO mice**: Mice lacking pyruvate dehydrogenase kinase 2 and 4 (double knock out; DKO), resulting in constitutively activated PDH, were kindly provided by Dr. Robert A. Harris.

**LC-MS/MS Analysis:** Plasma PK activator concentrations were determined by mass spectrometry using a SCIEX 5500 QTRAP equipped with a SelexION for differential mobility separation (DMS). Plasma samples were injected onto a C18 column (3μm particle size, 2.1x150 mm, Thermo Fisher Scientific) at a flow rate of 0.4 mL/min. Plasma sample were eluted with a combination of aqueous (A: 15mM ammonium formate and 10uM EDTA) and organic mobile phase (B: 60% acetonitrile, 35% isopropanol and 15mM ammonium formate) according to the following gradient: t=0min, B=0%; t=0.5min, B=0%, t=1min, B=40%; t=1.5min, B=40%; t=2min, B=0%; t=6min, B=0%. Metabolite detection was based on multiple reaction monitoring (MRM) in negative mode using the following source parameters: CUR: 30, CAD: high, IS: ±1500, TEM: 625, GS1: 50 and GS2: 55.

| Q1 | Q3 | t | ID | DP | EP | CE | CXP |
|---|---|---|---|---|---|---|---|
| 373 | 373 | 10 | Pka NIH 373/373 | 80 | 7 | 8 | 18 |
| 467 | 467 | 200 | PKa Agios 467/467 | -30 | -8 | -10 | -20 |
| 467 | 403 | 200 | PKa Agios 467/403 | -30 | -8 | -50 | -20 |

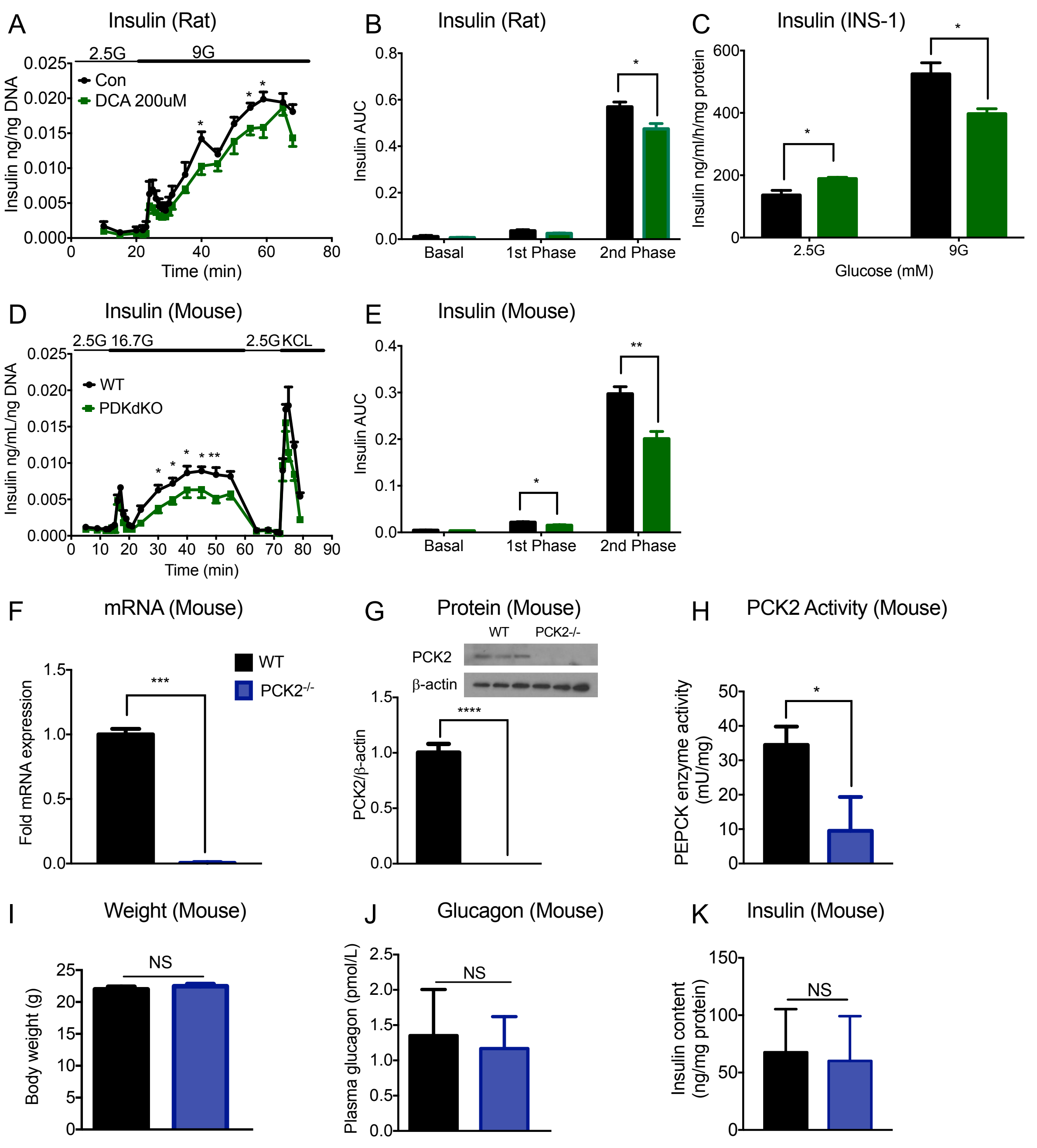


**Figure S1. Enhancing oxidative metabolism does not improve islet function and PCK2$^{-/-}$ mice have impaired beta-cell function in vivo.**

(**A- C**) Insulin secretion was measured in isolated islets from regular chow rats and INS-1 at the presence of PDK activator DCA (N=4). (**D and E)** First and second phase insulin secretion from perfused, isolated islets from control (WT), and PDKdKO mice. (**F**) PCK2 mRNA expression in isolated islets from WT and PCK2$^{-/-}$ mice. (**G**) PCK2 protein expression (N=3). (H) PCK2 activity was measured in islets from WT and PCK2$^{-/-}$ mice (N=4). (I) Body weight of WT and PCK2$^{-/-}$ mice. (**J**) Basal plasma glucagon level. (K) Total insulin content in islets of WT and PCK2$^{-/-}$ mice. Data are represented as mean ± SEM. of n=4-5 per group for PCK2 mRNA, PCK2 activity, total insulin content and 7-8 for body weight. Statistical comparisons made by t-test (*P<0.05, **P<0.01vs control wild type).

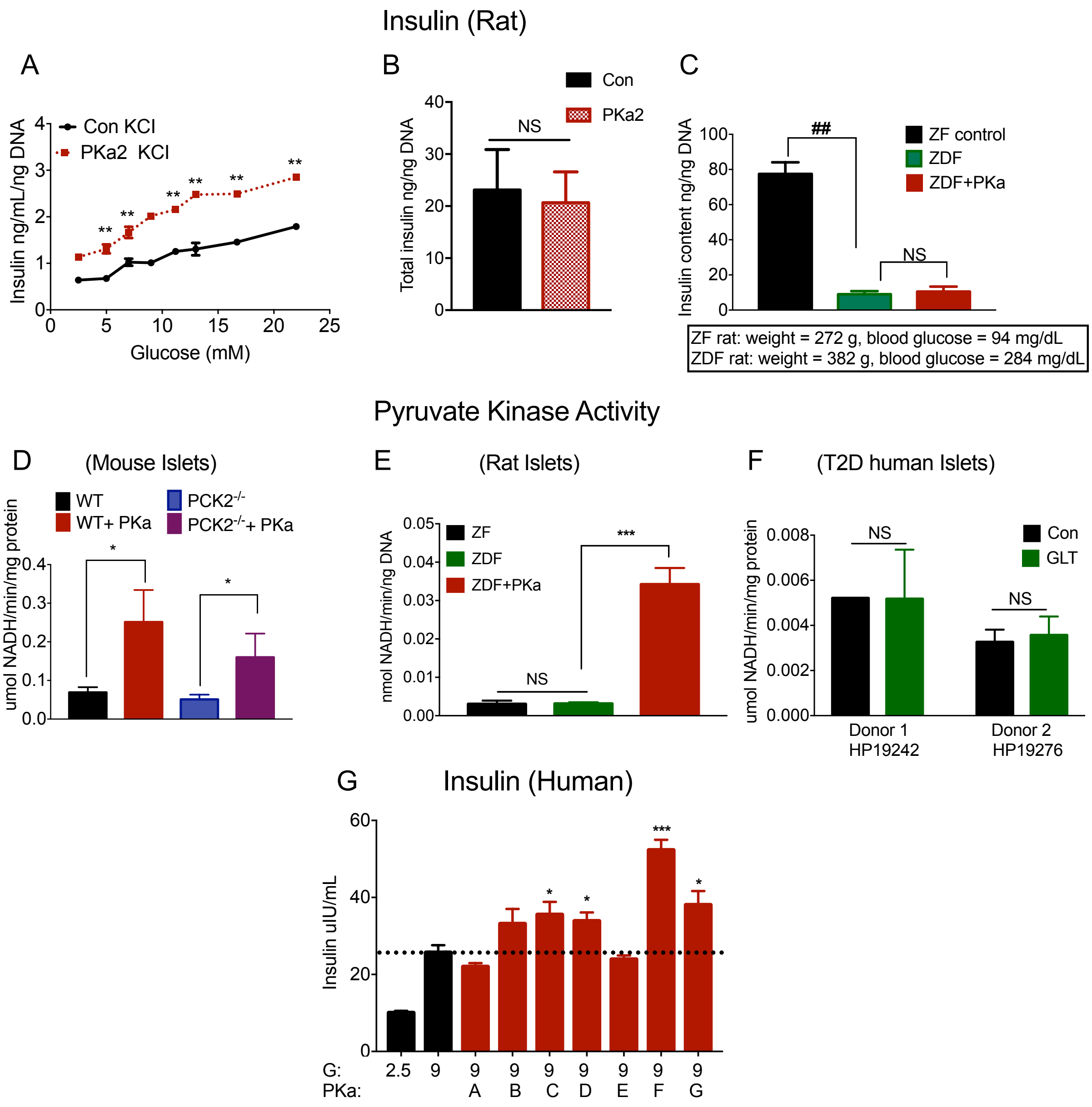


**Figure S2. PK activation improves insulin secretion in rodent and human models of T2DM.** **(A)** GSIS: Islet re-aggregates from HFD rats treated with PKa2 acutely at the condition of 30 mM KCL (n=4). **(B)** Islet total insulin content from HFD-fed rats chronically treated with PKa2 (n=4). **(C)** Total insulin content of islets from ZF, ZDF and ZDF rats treated with PKa (n=2). **(D)** PK activity was evaluated in islets from WT, $PCK2^{-/-}$ mice. **(E)** PK activity was confirmed in islets from ZF, ZDF and ZDF-diabetic rat treated with PKa. **(F)** PK activity of type 2 diabetic human islets. **(G)** PK activators increased the average insulin secretion of human islets in a static incubation in 9 mM glucose (9G). A, 10 µM NIH NCGC185916-06 (Dasa-58); B, 10 µM NIH NCGC188799-02; C, 10 µM TEPP-46; D, 10 µM NIH NCGC186527-05; E, 10 µM NIH NCGC181801-02; F, 10 µM NIH NCGC188795-01; G, 10 µM NIH NCGC183333-05. Data are represented as mean ± SEM of N=4-5. Statistical comparisons made by t-test and by 2-way ANOVA (*with vs. without PKa; $^{\#}$, ZF vs. ZDF or control vs. GLT; *,$^{\#}$, P<0.05; **,$^{\#\#}$, P<0.01).

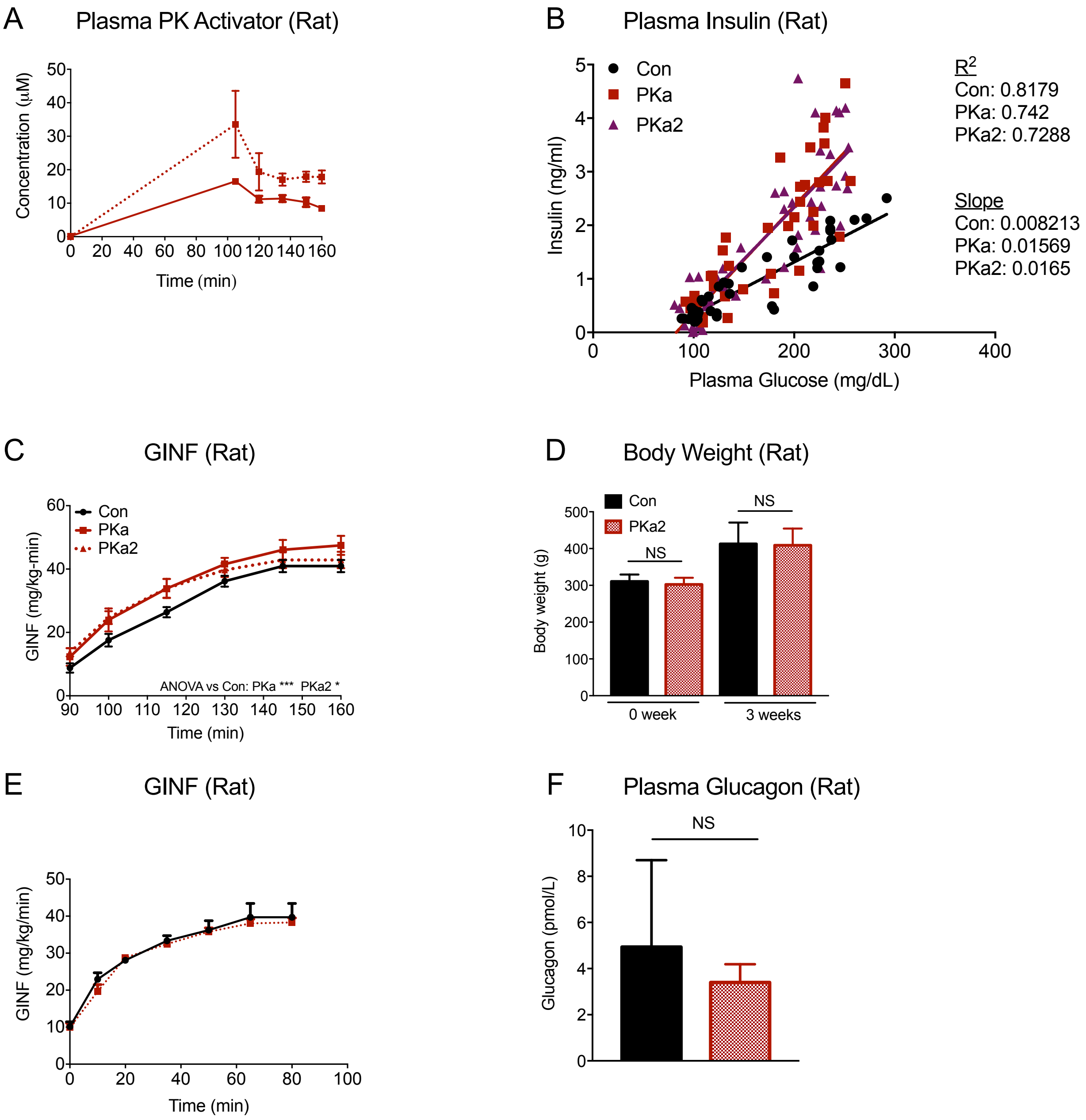


**Figure S3. PK activation improves insulin secretion and health *in vivo***

**(A)** Plasma concentration of PK activators after an infusion in chow-fed Sprague-Dawley rats. (**B**) The insulin vs glucose in hyperglycemic ramp of PK activators infused rats. (**C**) Glucose infusion rate to maintain hyperglycemia hyperglycemic ramp in rats acutely infused PK activators. (**D**) The body weight of rats before and after PKa2 chronic treatment. (**E**) Glucose infusion rate to maintain hyperglycemia during hyperglycemic ramp in HFD-fed rats chronically treated with PKa2. **(F)** Plasma glucagon levels of HFD-PKa2 and HFD-control. Data are represented as mean ± SEM of N=6-7. Statistical comparisons made t-test and by 2-way ANOVA (*, P<0.05, **, P<0.01).

**Table S1.** **Human islets donor characteristics related to figure 3.**

| Donor | Age (Years) | Sex | BMI | HbA1c (%) |
|---|---|---|---|---|
| H-108 | 32 | F | 36.2 | 5 |
| R-107 | 56 | F | 27.5 | 9.3 |
| HP 18038 | 45 | M | 27.3 | 6.5 |
| HP 18103 | 35 | F | 34 | 7.1 |
| HP 18212 | 40 | M | 27.7 | 7.4 |

**Table S2. Physiological amino acid mixture concentrations, Related to Figure 2**

| **Amino acid** | **Concentration at 1x (μM)** |
|---|---|
| Alanine | 2100 |
| Glutamine | 600 |
| Glycine | 700 |
| Valine | 550 |
| Leucine | 500 |
| Serine | 350 |
| Arginine | 200 |
| Lysine | 218 |
| Threonine | 121 |
